# Structural and functional connectivity reconstruction with CATO - A Connectivity Analysis TOolbox

**DOI:** 10.1101/2021.05.31.446012

**Authors:** Siemon C. de Lange, Koen Helwegen, Martijn P. van den Heuvel

## Abstract

We describe a Connectivity Analysis TOolbox (CATO) for the reconstruction of structural and functional brain connectivity based on diffusion weighted imaging and resting-state functional MRI data. CATO is a multimodal software package that enables researchers to run end-to-end reconstructions from MRI data to structural and functional connectome maps, customize their analyses and utilize various software packages to preprocess data. Structural and functional connectome maps can be reconstructed with respect to user-defined (sub)cortical atlases providing aligned connectivity matrices for integrative multimodal analyses. We outline the implementation and usage of the structural and functional processing pipelines in CATO. Performance was calibrated with respect to simulated diffusion weighted imaging from the ITC2015 challenge, test-retest diffusion weighted imaging data and resting-state functional MRI data from the Human Connectome Project. CATO is open-source software distributed under the MIT License and available as a MATLAB toolbox and as a stand-alone application at www.dutchconnectomelab.nl/CATO.

## Introduction

Network analysis of macroscale structural and functional brain connectivity data has become a widely used method in neuroscience (Bassett and Sporns, 2017). Neuroimaging data, including magnetic resonance imaging (MRI), can be used to make a reconstruction of the brain’s anatomical connections and functional synchronization patterns (Hagmann et al., 2008) and study the topological structure of the derived networks (Rubinov and Sporns, 2010). Network studies of the human and animal brain have led to the discovery of general organizational principles of healthy brain structure and function (Bullmore and Sporns, 2012) and abnormalities related to psychiatric and neurological disorders (Fornito et al., 2015).

MRI connectome studies aim to reconstruct and study structural and functional connectivity maps from diffusion weighted imaging (DWI) and resting-state functional MRI (rs-fMRI) data and the field is rapidly developing tools for the reconstruction of brain connectivity (for example (Aydogan and Shi, 2021; Cieslak et al., 2021; Cook et al., 2005; Cox and Hyde, 1997; Craddock et al., 2013; Esteban et al., 2019; Garyfallidis et al., 2014; Glasser et al., 2013; Jenkinson et al., 2012; Kiar et al., 2018; Leemans et al., 2009; Tourbier et al., 2022; Tournier et al., 2019; Wang et al., 2015; Wang et al., 2007; Whitfield-Gabrieli and Nieto-Castanon, 2012; Yeh, 2022) and see (Soares et al., 2013) and (Phinyomark et al., 2017) for reviews).These reconstruction tools and pipelines differ in their aims (e.g., the processed imaging modalities and included processing steps), programming language (e.g., C, MATLAB and python) and methodologies (e.g., probabilistic versus deterministic fiber tracking, data-driven versus predefined structural parcellations). The availability of multiple software packages is beneficial for researchers to be able to choose the software package suited for their study, expertise level and preferred programming framework.

We describe CATO – Connectivity Analysis TOolbox – a flexible toolbox for reconstructing structural and functional connectomes from DWI and rs-fMRI data. In the development of CATO, we put attention to the accessibility of CATO to users at all levels of expertise. For researchers starting out in connectomics, we provide documentation, both in this manuscript and on the documentation website (www.dutchconnectomelab.nl/CATO), together with an online configuration assistant that guides users through creating a configuration file for their specific dataset. For advanced users, CATO provides a configurable and modular structure with customizable pre-processing and the option to execute specific structural and functional processing steps. This flexibility allows users, for example, to choose FreeSurfer (Fischl, 2012), FSL (Jenkinson et al., 2012) and/or other tools for data preprocessing and to add their own processing steps to perform post-processing.

CATO allows for the reconstruction of structural and functional connectivity with respect to the same set of atlas templates. Matching connectivity reconstructions lets researchers investigate the interactions between both modalities, which can provide new insights into e.g. the dynamics of functional activity (Suárez et al., 2020) or the biological mechanisms of brain diseases (Cui et al., 2019; van den Heuvel et al., 2013). The structural pipeline in CATO offers diffusion reconstruction using diffusion tensor imaging (DTI), generalized q-sampling imaging (GQI) and constrained spherical deconvolution (CSD) and deterministic fiber tracking. CATO also has the feature to combine advanced diffusion reconstruction methods (GQI, CSD) in voxels with crossing fibers and simple and robust diffusion reconstruction (DTI) in voxels with fibers in a single direction. Functional connectivity is estimated using (partial) Pearson’s correlation coefficient.

CATO is developed in MATLAB and available as a MATLAB-based toolbox and as stand-alone application (that is also available in a Docker container). The implementation in MATLAB might be beneficial for researchers that are already familiar with the programming language, or who appreciate MATLAB’s efficient matrix algebra. The toolbox is open-source under the MIT License and publicly developed in the GitHub repository www.github.com/dutchconnectomelab/CATO.

In this paper we describe the steps in the structural and functional pipelines, followed by a description of the Configuration Assistant that provides a graphical user interface for creating and modifying configuration files for the separate pipelines. In the Results section, we benchmark and discuss performance with respect to the simulated diffusion dataset of the ISMRM 2015 Tractography challenge (ITC2015) (Maier-Hein et al., 2017) and functional data of the Human Connectome Project (Glasser et al., 2013).

## Methods

CATO is available as a package for MATLAB and as a compiled standalone executable for use on high-performance computer clusters. It has two separate pipelines for reconstructing structural and functional connectivity each consisting of multiple processing steps implemented as MATLAB functions (see Figure 1). The main functions `structural_pipeline` and `functional_pipeline` validate the user-provided input parameters and files and execute the desired pipeline steps. The preprocessing and parcellation functions utilize external software by executing a user-specified bash shell script (see Figure 2). The example preprocessing scripts and atlas parcellation scripts provided with CATO use FreeSurfer (Fischl et al., 2004) and FSL (Jenkinson et al., 2012) to perform data preprocessing and brain parcellation.

**Figure 1.**
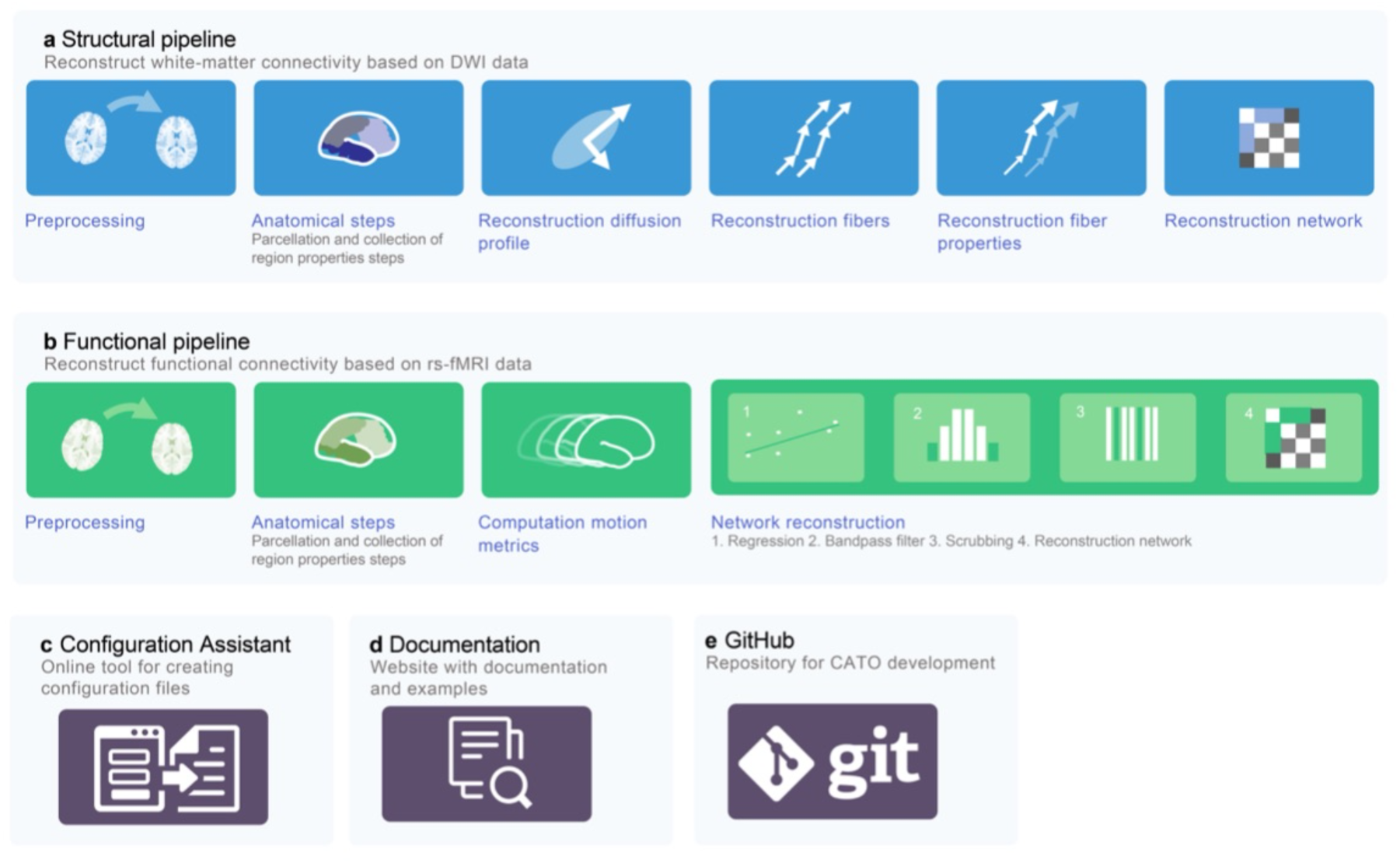
Overview CATO toolbox and pipelines. **a.** and **b.** CATO is a modular toolbox consisting of two pipelines for reconstruction of respectively structural (blue) and functional (green) connectivity. Each pipeline consists of multiple separate processing steps, with the anatomical parcellation steps being shared between pipelines. **c.** and **d.** The toolbox is accompanied by an online configuration assistant that helps users generate configuration files and online documentation. **e.** The CATO toolbox is developed on a public GitHub repository.

**Figure 2.**
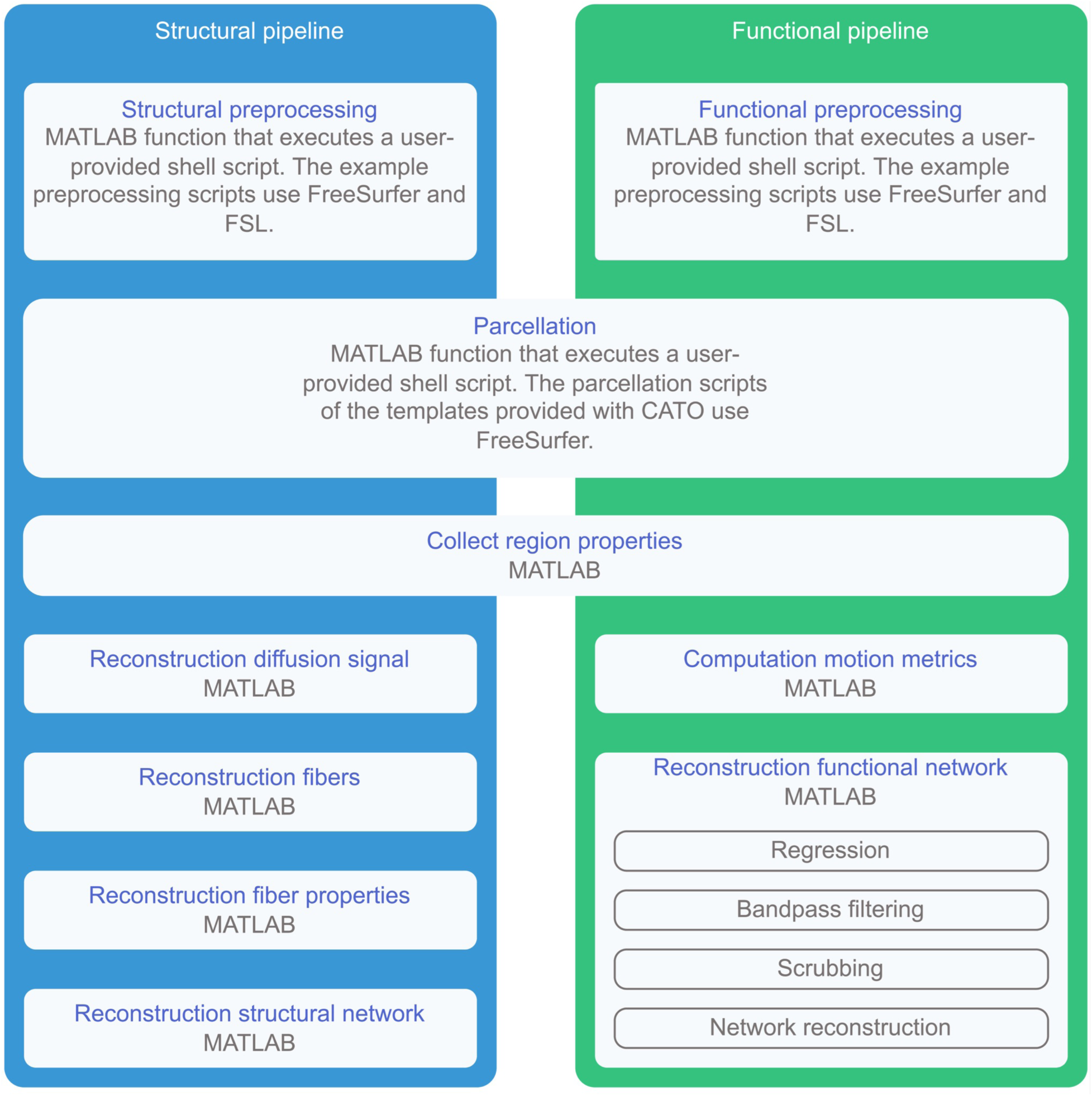
Implementation overview of the structural and functional pipeline and its dependencies. Overview of structural and functional pipeline steps and the programming language used (MATLAB or MATLAB and shell code). The parcellation and ‘collect region properties’ steps are shared between the structural and functional pipelines. External software can be used in the user-provided bash shell scripts that are executed in the structural preprocessing, functional preprocessing and parcellation step. The example scripts provided with CATO use FreeSurfer (preprocessing and parcellation steps) and FSL (preprocessing step).

### Part I: Structural pipeline

Reconstruction parameters for the structural pipeline can be specified on the command line or in a configuration file in the JSON-format. The reconstruction parameters describe e.g. the desired processing steps (detailed below), location of the anatomical FreeSurfer directory, location of the DWI files, location of the gradient information files, cortical atlas types, and settings for preprocessing, diffusion reconstruction, fiber tracking and network reconstruction.

#### Structural preprocessing

The structural pipeline requires preprocessed T1 data (for tissue classification and cortical parcellation) and a DWI set (we recommend 30 or more diffusion directions (Jones, 2004)). We recommend the automatic FreeSurfer cortical reconstruction process to preprocess structural T1 data (Fischl et al., 2004). Performing FreeSurfer cortical reconstruction prior to running CATO provides the required files needed for downstream network reconstruction and improves the computational efficiency when processing subjects in parallel. For the DWI data, we recommend the use of the elaborate FSL package for preprocessing of the DWI data (Jenkinson et al., 2012). Required structural preprocessing steps can be executed by a bash shell script specified by the parameter `preprocessingScript` allowing users to customize a preprocessing script that fits best to their data, or exclude preprocessing all together and start with already pre-processed data. Three example preprocessing scripts (FSL topup-eddy-preprocessing, FSL eddy-preprocessing and minimal-preprocessing) are provided with the structural pipeline that perform all needed steps and data output files for subsequent fiber tracking and network reconstruction. These example preprocessing scripts call the well-documented and thoroughly evaluated FSL `topup` and FSL `eddy` tools to correct for susceptibility induced distortions, eddy current distortions and motion artifacts in the DWI data (Andersson and Sotiropoulos, 2016), update the b-vectors to adjust for DWI corrections (Leemans and Jones, 2009), compute a DWI reference image based on the corrected diffusion-unweighted (b0) volumes, compute the registration matrix between DWI reference image and the anatomical T1 image using FSL’s `bbregister` (Greve and Fischl, 2009) and registers the FreeSurfer segmentation to the DWI reference image.

#### Anatomical parcellation – Parcellation

`Parcellation` creates cortical parcellations of the surface with respect to reference atlases. Atlases included with CATO are the Desikan-Killiany atlas present in FreeSurfer (Desikan et al., 2006), combined with the 120, 250 and 500 regions Cammoun sub-parcellations of the Desikan-Killiany atlas (Cammoun et al., 2012) and the Von Economo-Koskinas cortical region and cortical-type atlas (Scholtens et al., 2016). Each reference atlas has a directory within the `TOOLBOXDIR/templates` directory containing a parcellation script (bash script), an ROIs file and template-specific files. File names are defined as variables (`templatesDir`, `templateScript`, `ROIsFile`) in the configuration file and can be modified by the user. The user can add surface-based or volume-based atlases by adding a new directory with template-specific files to the CATO templates directory. During the parcellation step, the parcellation script (`templateScript`) is executed such that the brain surface and volume is parcellated with respect to the reference atlas. The parcellation scripts provided with CATO use FreeSurfer to: 1. create annotation files for the atlases in the subject’s FreeSurfer directory (using `mris_ca_label`). 2. Map the new cortical labels to the automated segmentation (FreeSurfer aseg) file in the subject’s FreeSurfer directory (using `mri_aparc2aseg`). 3. Map this combined volume to the DWI reference image creating the subject’s `parcellationFile` in the output directory of CATO (using `mri_label2vol`). 4. Create anatomical statistics files in the subject’s FreeSurfer directory containing for each region, among others, the number of vertices, surface area (mm^2^), gray matter volume (mm^3^) and average thickness (mm) (using `mris_anatomical_stats`).

#### Anatomical parcellation – Summarized region properties

The `collect_region_properties` step collects volumetric and surface data of brain regions and summarizes this data in the `regionPropertiesFile` (MATLAB-file) The following statistics are included for every brain region: *i)* center of mass of each region (calculated from the parcellation file), *ii)* the number of vertices, *iii)* surface area (mm^2^), *iv)* gray matter volume (mm^3^), and *v)* average thickness (mm) (all from FreeSurfer’s stats file). These metrics are used in later steps of the pipeline.

#### Reconstructed diffusion signal

The `reconstruction_diffusion` step estimates the white matter fiber organization in each voxel from the measured DWI data. Structural connectivity modeling is based on the principle that white matter fibers restrict the movement of water molecules resulting in peaks (preferred diffusion directions) in the diffusion signal in voxels (Mori et al., 1999). The `reconstruction_diffusion` step provides three methods to infer the diffusion peaks from DWI data, including Diffusion Tensor Imaging (DTI), Constrained Spherical Deconvolution (CSD) and Generalized Q-sampling imaging (GQI) and the user can implement other methods when required. For each method, reconstruction parameters are provided in the configuration file. For each method, the reconstructed peaks of all voxels are saved in the `diffusionPeaksFile` (MATLAB-file) and associated diffusion measures are saved in the `diffusionMeasuresFile` (MATLAB-file).

#### Diffusion Tensor Imaging (DTI)

`DTI` reconstruction models the signal measured in a voxel by a single tensor that defines the diffusion profile associated with one preferred diffusion-direction. Tensor estimation is performed using the informed RESTORE algorithm (Chang et al., 2005) that performs tensor estimation while identifying and removing outliers during the fitting procedure, reducing the impact of physiological noise artifacts on the diffusion tensor modeling (de Reus, 2015; Haddad et al., 2019) (see Supplementary Methods 1). The Levenberg-Marquardt method (based on implementation by Gavin (Gavin, 2019) is used to solve the nonlinear least squares problem (see Supplementary Methods 1). Four diffusivity measures are computed from the estimated tensor, including: fractional anisotropy, axial diffusivity, radial diffusivity and mean diffusivity are computed (Alexander et al., 2007; Basser and Pierpaoli, 1996).

#### Constrained Spherical Deconvolution (CSD)

`CSD` reconstructs the fiber orientation distribution function (fODF) in a voxel by deconvolution of the measured signal with the diffusion profile associated with a fiber (Tournier et al., 2007). Signal deconvolution is performed in the super-resolution spherical harmonics framework to allow for a natural description of the surface of a diffusion process (the used order of spherical harmonics is set by parameter `shOrder`). The spherical deconvolution is constrained to non-negative spherical harmonics which reduces high frequency noise and results in the reconstruction of a few well-defined peaks (Tournier et al., 2007). Parameter `lambda` specifies the regularization parameter λ that controls the coarseness of the reconstructed fODF. Parameter `tau` specifies the amplitude threshold τ below which the corresponding fODF is assumed zero. The implementation is based on the CSD reconstruction method in the Dipy software package (Garyfallidis et al., 2014) and as described in Tournier et al. (2007). Diffusion peaks are selected from the local maxima of the reconstructed fODFs. Local maxima are considered diffusion peaks if their fODF value, normalized by the maximum value of the fODF, was larger or equal to the `minPeakRatio`. The maximum number of largest local maximas that are selected as diffusion peaks is set by the `outputPeaks` parameter. Voxels with more than `maxPeaks` number of largest local maximas are considered to have an isotropic diffusion profile and no diffusion peaks were selected.

#### Generalized Q-sampling Imaging (GQI)

`GQI` reconstructs the fODF in a voxel by utilizing the Fourier transform relation between the diffusion signal and the underlying diffusion displacement (Yeh et al., 2010). The implementation follows the method described in Yeh et al. (Yeh et al., 2010) and the example code as presented on http://dsi-studio.labsolver.org. Parameter `meanDiffusionDistanceRatio` specifies the mean diffusion distance ratio that regulates the coarseness of the fODF. Diffusion peaks were reconstructed from the fODF using the same method as described above for the CSD reconstruction method. Diffusion measures generalized fractional anisotropy (Tuch, 2004) and quantitative anisotropy (Yeh et al., 2010) of each peak are also calculated and added to the `diffusionMeasuresFile`.

#### Combined implementation

Comparing the three reconstruction methods, DTI is a robust and relatively simple method, with CSD and GQI being more advanced diffusion models that allow a better description of the diffusion signal in voxels having a more complex underlying white matter organization. To combine the strengths of simple and advanced reconstruction methods, CATO gives users the option to combine DTI with CSD or GQI. This option performs DTI modeling (being more robust) in voxels where CSD or GQI estimate only one peak, and uses CSD or GQI in voxels where multiple peaks are detected (being more flexible in voxels with a complex fiber architecture). Specific parameter settings for each reconstruction method are set in the configuration file and are described in more detail in the online documentation.

#### Intermediate output files

The `reconstruction_diffusion` step provides the user the additional option to export diffusion measures to a NIFTI volume file `diffusionMeasuresFileNifti`. Standard diffusion measures that are exported include fractional anisotropy, axial diffusivity, radial diffusivity and mean diffusivity.

#### Reconstructed fibers

The `reconstruction_fibers` step performs fiber tracking based on the diffusion peaks in each voxel. The fiber tracking step utilizes the standard “Fiber Assignment by Continuous Tracking” (FACT) algorithm (Mori et al., 1999) (adjusted for multiple peaks when CSD, GQI or combined methods are used). FACT describes a deterministic tracking algorithm that starts fiber reconstruction from seeds in the white matter and propagates streamlines in the main diffusion axis of the voxel while updating the propagation direction each time the tip of the streamline enters a new voxel. The CATO implementation of FACT starts from one or multiple seeds (number of seeds set by `NumberOfSeedsPerVoxel`) in all voxels with a segmentation label matching the list provided by `startRegions`. Fiber reconstruction stops if a tracker is *i)* in a region with low fractional anisotropy (FA < `minFA)`, *ii)* in a stopping region with segmentation code included in the `stopRegions` variable, *iii)* about to revisit the current voxel or previously visited voxel, *iv)* about to enter a forbidden region with segmentation code included in the `forbiddenRegions` variable or *v)* about to make a sharp turn (with angle > `maximumAngleDeg`).

#### Reconstructed fiber properties

The `reconstruction_fiber_properties` step identifies fiber segments that connect brain regions and calculates fiber measures as preparation for the network reconstruction step. For each atlas and reconstruction method, the `reconstruction_fiber_properties` step iterates through all fibers and if a fiber crosses two or more brain regions of interest, as defined by the regions of interest file (`ROIsFile`, text-file), then the shortest fiber segment between each region pair is included in the fiber properties file (`fiberPropertiesFile`, MATLAB-file). In addition to the start and end point of each fiber segment and the associated region pair, additional measures for fiber segments are stored, including: *maximum turn angle, minimum fractional anisotropy, fiber length* (physical length in mm) and the average *fractional anisotropy*, *axial diffusivity*, *radial diffusivity*, *mean diffusivity* and *generalized fractional anisotropy.* Diffusion measures are averaged over voxels weighted by the length of the traversed path through each voxel (Tuch, 2004).

#### Structural connectivity matrix

The `reconstruction_network` step builds the connectivity matrices for each of the selected (sub)cortical atlases and reconstruction methods. Brain regions included in the connectivity matrix, and their order, are defined by the regions of interest file (`ROIsFile`). The network matrix is constructed by iterating through all fiber segments in the `fiberPropertiesFile` and fiber segments connecting regions of interest are added to the connectivity matrix.

In addition to the fiber reconstruction criteria used in the `reconstruction_fibers` step, fibers can additionally be filtered in the `reconstruction_network` step by their projection length (with only fibers longer than `minLengthMM` being included in the network reconstruction), minimum fractional anisotropy (including only fibers that touch voxels with fractional anisotropy higher than `minFA`) and maximum angle (including only fibers that make turns in their trajectories smaller than `maxAngleDeg`). Connectivity matrices are saved to the file `connectivityMatrixFile` (MATLAB-file). Connectivity matrices include weights by the *number of streamlines*(i.e. the number of fibers) that connect two regions, *fiber length* (physical length in mm averaged across fibers), the average *fractional anisotropy*, *axial diffusivity*, *radial diffusivity*, *mean diffusivity*, *generalized fractional anisotropy, streamline volume density* and *streamline surface density* (Tuch, 2004) of the reconstructed streamline segments.

### Part II: Functional connectivity

The functional connectivity pipeline builds upon the same modular organization as the structural connectivity pipeline, including a preprocessing step, two anatomical steps and a connectivity reconstruction step (see Figure 1). Resting-state fMRI processing starts with the main function `functional_pipeline` that runs the processing steps according to the parameters provided on the command line or rs-fMRI pipeline configuration file (JSON-file). Reconstruction parameters describe e.g. the desired processing steps (detailed below), location of the anatomical FreeSurfer directory, location of the rs-fMRI file, cortical atlas types, and settings for preprocessing, regression, bandpass filtering, scrubbing and network reconstruction. The functional pipeline includes the following steps:

#### Preprocessing

The step `functional_preprocessing` provides users the option to preprocess rs-fMRI data with their preferred methods, specified in a bash script (`preprocessingScript`). The default preprocessing script performs steps for correcting slice timing using FSL tool `slicetimer` and correcting motion artifacts using FSL tool `MCFLIRT` (Jenkinson et al., 2002), computing a rs-fMRI reference image by averaging all (motion corrected) rs-fMRI frames (using FSL tools), computing a registration matrix between the rs-fMRI reference image and the T1 image (Greve and Fischl, 2009) (using FreeSurfer) and registration of the T1 parcellation to the rs-fMRI image (using FreeSurfer). The modular pipeline allows preprocessing with other packages and completely custom preprocessing when required. Additional example preprocessing scripts, for e.g. BIDS-organized data and preprocessing of rs-fMRI data using ICA+FIX, are provided on the documentation website.

#### Anatomical parcellation

The first steps of the functional pipeline include the same parcellation steps `parcellation` and `collect_region_properties` as the structural pipeline to parcellate the cortical surface with respect to the same atlases as the structural pipeline, such that structural-functional relationships can be explored in post-processing integrative multimodal analyses. The anatomical steps further compute the anatomical statistics of these regions and collect the region properties into the region properties file `regionPropertiesFile` (MATLAB-file).

#### Computed motion metrics

`Compute_motion_metrics` computes for each frame the motion metric framewise displacement (FD) and the change in signal intensity between frames (known as ‘DVARS’) (Power et al., 2012). Measures are derived from a `motionParametersFile` file that can be created in the preprocessing step with MCFLIRT (Jenkinson et al., 2002) and are saved in the `motionMetricsFile` (MATLAB-file). Framewise displacement (FD) is computed as the sum of the estimated translational and rotational displacement in a frame (Power et al., 2012), with rotational displacement defined in degrees in the `motionParametersFile` file (MATLAB-file). FD is converted to millimeters by calculating the expected displacement on the surface of a sphere of radius 50 mm as model for the cerebral cortex (Power et al., 2012). DVARS is calculated as the square root of the average squared intensity differences of all brain voxels between two consecutive rs-fMRI frames (Power et al., 2012).

#### Functional connectivity matrix

The `reconstruction_functional_network` step computes the functional connectivity matrices preceded by (optional) covariate regression, bandpass filtering and scrubbing of the time-series.

#### Regression

Per voxel, covariates (referred to as regressors) are removed from the rs-fMRI time series by calculating the residuals of a linear model of the signal intensities with the regressors as predictors. Standard regressors include the linear trends of motion parameters, first order drifts of motion parameters and mean signal intensity of voxels in white matter and CSF (specific regions are defined by configuration parameter `regressionMask` and each region code in the regression mask is included as separate regressor). Mean signal intensity of all voxels in the brain can optionally be included as an additional regressor to perform global mean correction (indicated by `globalMeanRegression`) (Fox et al., 2009).

#### Bandpass filtering

Next, (optional, but recommended) bandpass filtering to the rs-fMRI data is applied (indicated by parameter `filter`). A zero-phase Butterworth bandpass filter is used with low-pass and high-pass cutoff frequencies set by the configuration parameter `frequencies`.

#### Scrubbing

Frames that display significant motion artifacts are removed from the rs-fMRI time-series (scrubbing) (Power et al., 2012). When scrubbing is applied (optional, but highly recommended), frames with motion artifacts are identified based on two indicators: *i)* having framewise displacement FD larger than `maxFD` and *ii)* having a DVARS larger than Q3 + `maxDVARS` × IQR, where IQR refers to the interquartile range IQR = Q3 – Q1, with Q1 and Q3 referring to the first and third quartile of the DVARS of all frames. Frames with a number of indicators larger or equal to `minViolations` are labeled as frames with potential motion artifacts and are excluded from further analysis. To accommodate temporal smoothing of data, frames consecutive to frames with labeled motion artifacts are optionally excluded: configuration parameter `backwardNeighbors` determines the number of preceding frames and `forwardNeighbors` determines the number of succeeding frames to be excluded from further analysis.

#### Network reconstruction

The `reconstruction_functional_network` step computes region-to-region functional connectivity. Functional connectivity is estimated between brain regions (specified by the `ROIsFile`) as the correlation coefficient of the average signal intensity of these regions across the selected frames. The standard correlation measure (specified by `reconstructionMethod`) is the Pearson’s correlation coefficient. Connectivity matrices are saved to the file `connectivityMatrixFile` (MATLAB-file) and time series of each region are saved to the file `timeSeriesFile` (MATLAB-file). Multiple `reconstruction_functional_network` steps can be performed to explore different reconstruction settings and are distinguished by using the parameter `methodDescription`.

### Configuration assistant

The Configuration assistant is an online tool to create and modify configuration files for the structural and functional pipeline (http://www.dutchconnectomelab.nl/CATO/configuration-assistant). The Configuration assistant checks all configuration parameters for validity.

### Development

CATO is publicly developed in the GitHub repository https://github.com/dutchconnectomelab/CATO. This repository provides a history of software releases, current development and automated software tests. Users can use this repository to submit feature requests, report bugs or contribute to the software. Development principles, including contribution flow, style guides and code testing, are provided in the contribution guidelines of the repository.

### Structural pipeline ground-truth benchmarking

#### Data

The variety of implemented reconstruction methods enables researchers to select a preferred trade-off between sensitivity and specificity in connectome reconstructions (Zalesky et al., 2016). We compared the used diffusion modeling and fiber reconstruction methodology with data of the ISMRM 2015 Tractography challenge (ITC2015) (Maier-Hein et al., 2017) to benchmark CATO to other reconstruction strategies and to evaluate the impact of different diffusion modeling methods and parameters. ITC2015 allows tractography pipelines to be evaluated with respect to a defined ‘ground truth’ white matter skeleton of 25 distinct white matter bundles distilled from DWI data of a single Human Connectome Project (HCP) subject by an expert radiologist (Glasser et al., 2013), including cortico-cortical, cortico-spinal and cortico-cerebellar tracts (see Supplementary Table 1 for a description of the included white matter bundles and Figure 3). This predefined ITC2015 white matter skeleton was used to subsequently simulate DWI images, which were distributed across research groups for analysis. Reconstructed fiber clouds from each research group were collected in the ITC2015 competition for benchmarking. We reconstructed four fiber clouds (for the method DTI, CSD, GQI, CSD-DTI and GQI-DTI combined) using CATO (version 3.2.0) with default HCP reconstruction parameters and a series of fiber clouds in which reconstruction parameters were adjusted (see Supplementary Methods 2 for a detailed description of the applied reconstruction steps and reconstruction parameters). The reconstructions were benchmarked on two key reconstruction features, including 1) the correctness of the spatial trajectory of reconstructed white matter bundles, and 2) the specificity and sensitivity of the reconstructed connectome networks.

**Figure 3.**
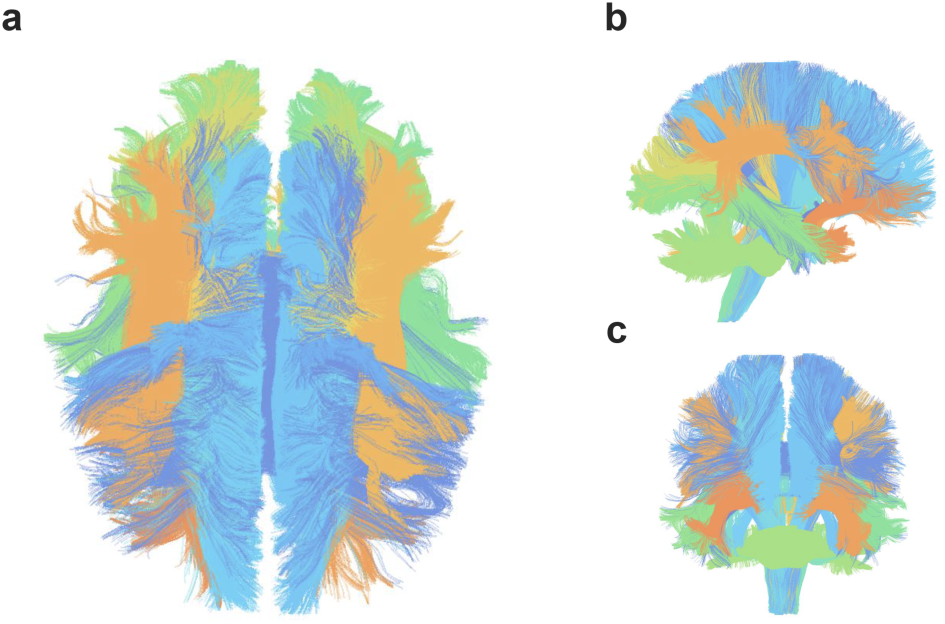
Ground truth white matter skeleton of the ISMRM 2015 Tractography challenge. a. Top view b. left-side and c. front.

#### Spatial reconstruction measures

We evaluated the spatial trajectory of the reconstructed fiber clouds with respect to three quality metrics using scripts provided by ITC2015 (Maier-Hein et al., 2017). The first two metrics are based on clustering fibers into white matter bundles and subsequently computation of their spatial overlap with the ITC2015 ground truth white matter bundles. This provides 1) the number of true-positive reconstructed white matter bundles (i.e. number of valid bundles, VB), 2) the number of reconstructed false-positive bundles (i.e. number of invalid bundles, IB). 3) The third metric counts the number of fibers with trajectories that spatially overlap with the ground truth white matter skeleton (i.e. number of valid fibers, VF). The ITC2015 referred to this third measure as ‘valid connections’; we here refer to it as ‘valid fibers’ to avoid confusion with connectome reconstruction measures.

#### Connectome reconstruction measures

We also evaluated the quality of the connectome reconstructions (a primary aim of CATO) using the ITC2015 data. To this end, fiber clouds from all ITC2015 submissions were downloaded from the data archive on Zenodo (https://doi.org/10.5281/zenodo.840086) and converted to 87 fiber cloud files in TRK-format using Dipy resulting (9 submissions were excluded due to errors including 15_0, 7_1, 7_2, 7_3, 3_3, 6_0, 6_1, 6_2 and 6_4). The anatomical T1-weighted image of the ITC2015 was parcellated into 68 distinct cortical regions according to the FreeSurfer Desikan-Killiany atlas (describing 34 left-hemispheric and 34 right-hemispheric cortical regions, (Desikan et al., 2006), using FreeSurfer version 6.0.0 (Fischl et al., 2004)). Using this parcellation, we computed 1) a ground truth connectome based on the ITC2015 ground truth white matter tracts, 2) connectome maps based on the fiber clouds from all ITC2015 submissions and 3) connectome maps based on the fiber clouds obtained from CATO reconstructions of the ITC2015 dataset (see Supplemental Methods 1 for details on the CATO reconstruction of the ITC2015 data). Fibers in the ground truth fiber cloud and fiber clouds from ITC2015 submissions were extended, at most two centimeters, to ensure that they all reach, on both fiber ends, non-white matter voxels (see Supplementary Figure 1). All three series of connectome maps were formed by combining the cortical parcellation with the respective fiber clouds into connectivity matrices using the `reconstruction_fiber_properties` and `reconstruction_network` functions, thus using identical processing steps for all connectome reconstructions. Connectome reconstruction quality was then assessed by means of computing the false positive rate (100% × number of false positive connections / number of absent connections in ground truth connectome) and false negative rate (100% × number of false negative connections / number of connections in ground truth connectome) (Zalesky et al., 2016) as measured with respect to the ITC2015 ground truth connectome.

### Structural and functional pipeline test-retest benchmarking

Test-retest reliability of connectome reconstructions was examined using the HCP test-retest cohort of 45 subjects (Glasser et al., 2013). For this subset of HCP subjects imaging scans are available of two visits separated by 18 to 343 days. We analyzed structural, functional and diffusion data that was preprocessed by HCP using the established HCP preprocessing pipeline that includes spatial artifact and distortion removal, cross-modal image registration and FreeSurfer reconstruction. Structural and functional connectomes were reconstructed using CATO (version 3.2.0) (see Supplementary Methods 3 for a detailed description of the applied reconstruction steps and reconstruction parameters). The measured test-retest reliability includes biological variance (aging effects) and methodological variance including aspects related to the data acquisition (e.g. scanner effects) and variance induced in the data processing. Test-retest benchmarking results are intended to inform on what level of test-retest reliability is to be expected when using CATO processed connectomes.

Test-retest reliability of connectivity strength was estimated by intra-class correlation in line with previous studies (Elliott et al., 2019; Tozzi et al., 2020). The ‘C-1’ intra-class correlation (ICC), also known as the ICC (3,1) (Shrout and Fleiss, 1979), was calculated for each connection across visits as

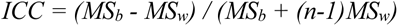

to quantify the degree of absolute agreement among scans across subjects, with *MS_b_* the between-subjects variances estimated by the between-subject mean squares of connection strength, *MS_w_* the within-subject variances estimated by the within-subject mean squares of connection strength and *n* the number of scans (*n*= 2) per subject (McGraw and Wong, 1996). The ICC implementation (ICC_C_1 from https://github.com/jmgirard/mReliability) corrected the ICC measure for missing data of subjects in which a connection was not reconstructed. Reliability was interpreted using the description by Cicchetti (Cicchetti, 1994) stating that ICC < 0.4 reflects poor reliability, ICC = [0.40 - 0.59] reflects fair reliability, ICC = [0.60 - 0.74] reflects good reliability and ICC = [0.75 - 1.00] reflects excellent reliability.

For structural connectivity benchmarking, test-retest reliability was calculated for the number of streamlines (NOS) and fractional anisotropy (FA) connectivity strength measures. We selected for each reconstruction method the top 26.6% connections with the highest prevalence across subjects to have a fair comparison between reconstruction methods (de Reus and van den Heuvel, 2013). This threshold was chosen such that the filtered connections of the GQI-DTI reconstruction were present in at least 60% of the subjects (de Reus and van den Heuvel, 2013). In the analyses across atlases, the number of connections differed between atlases and connections were selected such that they were present in 60% of the subjects. The ICC of connections was evaluated for four common cortical atlases (Desikan-Killiany atlas, the 120 and 250 subdivisions of the Desikan-Killiany atlas and the Von Economo-Koskinas atlas) and five reconstruction methods (DTI, GQI, CSD, GQI-DTI, CSD-DTI).

For the functional test-retest reliability experiment, connections were selected by a proportional threshold that selected 20% strongest correlations in a group-averaged functional connectome obtained by averaging connectivity matrices across subjects and visits. The ICC of connections was evaluated for the same four cortical atlases as in the structural test-retest analysis. The effect of global mean correction, bandpass filtering and scrubbing were also explored by evaluating ICC scores associated with the application of these reconstruction options. The test-retest reliability of the full matrix is presented in Supplementary Figure 2 and Supplementary Tables 2 and 3.

## Results

### Structural pipeline benchmarking

#### Spatial reconstruction

Performance was validated in comparison with tractography-based fiber clouds from (in total) 96 submissions of 20 different sites of the ITC2015 challenge. ITC2015 submissions showed a wide range of scores on the spatial reconstruction quality measures: the number of validly reconstructed bundles ranged from 5 to 24 bundles out of the 25 ground-truth bundles and the number of invalid bundles ranged from 27 to 386 bundles (Figure 4a). In reconstructions obtained with CATO 23 of the 25 ground-truth bundles (92%) were reconstructed, which was similar to the median of 23 of the ITC2015 submissions (Figure 4a). White matter bundles that were missed included the anterior commissure and posterior commissure. Across reconstruction methods, 62 to 73 invalid bundles were traced, which were not included in the ground-truth white matter skeleton (CSD: 73, DTI: 69, GQI: 62, GQI-DTI: 69 and CSD-DTI: 81), numbers that are comparable to ITC2015 submissions which had a median value of 77 invalid bundles (Figure 4a). The percentage of fibers with tractographies that spatially overlapped with the ground truth white matter skeleton ranged between 3.75% and 92.5% for the ITC2015 submissions (median: 62.6%), with CATO values ranging between 43% and 65% (CSD: 43%, DTI: 65%, GQI: 53%, GQI-DTI: 65% and CSD-DTI: 48%, Figure 4b).

**Figure 4.**
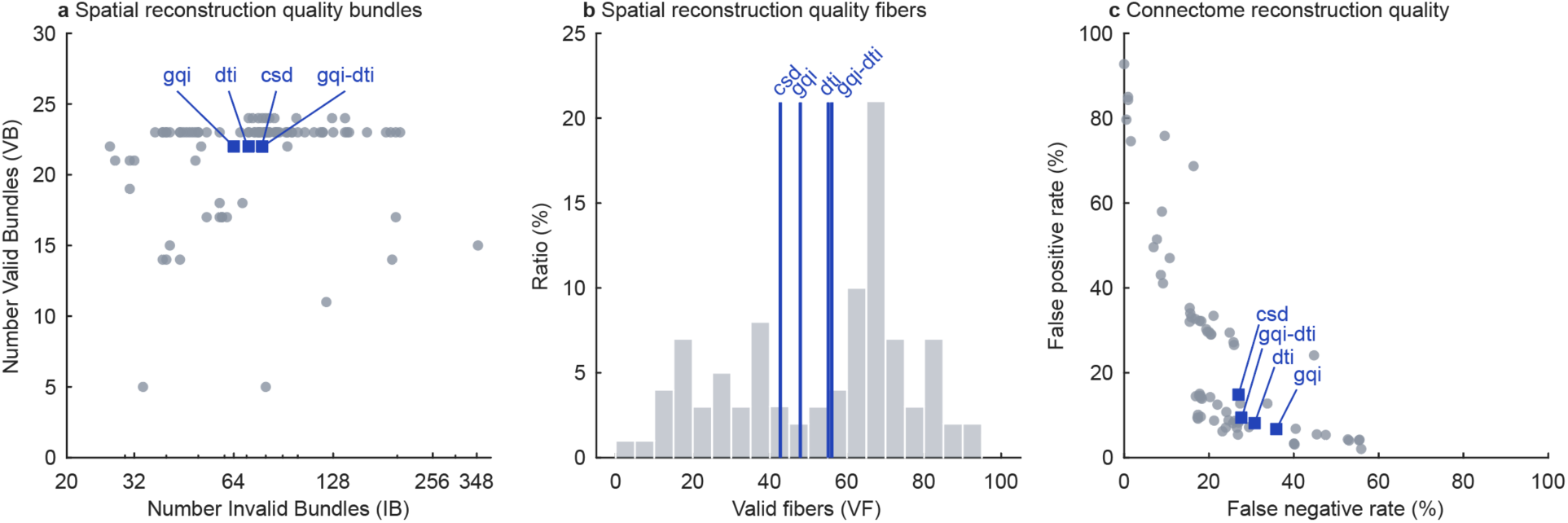
Benchmarking reconstructions. The spatial reconstruction quality of fiber cloud reconstructions obtained using CATO were benchmarked with respect to different reconstruction settings, and submissions in the ITC2015 challenge. **a.** The spatial reconstruction quality of fibers was assessed by the number of valid and invalid reconstructed bundles. The results of CATO are indicated by blue squares and results from submissions to the ITC2015 are indicated by gray circles. Reconstructions showed a similar number of valid bundles (23) and invalid bundles (62-73) as ITC2015 submissions (median VB = 23, median IB = 77). **b.** The spatial reconstruction quality was also assessed by the percentage of valid reconstructed fibers. The results of CATO are presented by blue lines and the gray histogram shows the valid fiber distribution across ITC2015 submissions. The percentage of valid fibers in the reconstructions ranged between 43% and 65% and ITC2015 submissions showed percentages between 3.75% and 92.5% (median VF = 62.6%). **c.** Connectome reconstruction quality (aim of CATO) was assessed by the level of false negative and false positive connection reconstructions. CATO showed relatively high specificity (false positive rates: 4% - 12%; and false negative rates: 29% - 43%) compared to the ITC2015 submissions (median false positive rates: 14.8%; and median false negative rates: 26.8%).

#### Connectome mapping

ITC2015 submissions (67 submissions included) showed a wide range of results, with false positive rates between 3% - 92% (median = 14.8%) and false negative rates between 0% - 69% (median = 26.8%). This emphasizes a wide spectrum of specificity-sensitivity trade-offs across the different strategies used by groups participating in ITC2015. CATO connectome reconstructions included false positive rates between 4% - 12% (CSD: 12%, DTI: 7%, GQI: 4%, GQI-DTI: 5% and CSD-DTI: 13%) and false negative rates between 29% - 43% (CSD: 29%, DTI: 32%, GQI: 43%, GQI-DTI: 36% and CSD-DTI: 26%, see Figure 4c). This relatively low false positive rate places CATO reconstructions at default settings as a connectome reconstruction tool with a relative conservative reconstruction sensitivity and relatively high connectivity reconstruction specificity (Zalesky et al., 2016). Levels of sensitivity and specificity of connectome reconstruction can further be tuned by varying the used reconstruction parameters, with diffusion reconstruction parameters showing small impact and fiber reconstruction parameters having large impact on the sensitivity-specificity trade-off (see Supplementary Results 1).

### Structural test-retest benchmarking

Test-retest reliability of the reconstructed structural connectivity was estimated in the HCP test-retest dataset. Reliability of connections was assessed by ICC, across various cortical parcellations, reconstruction methods and connectivity strength measures. The number of streamlines (NOS) connectivity strength measure showed, in the group-masked connectomes, in 19% of the connections ‘poor’, in 24% ‘fair’ and in 57% ‘good’ to ‘excellent’ reliability median (ICC=0.66, IQR=0.47-0.79, Desikan-Killiany atlas, GQI-DTI method). The median ICC varied between 0.59 - 0.70 across atlases with percentages of connections with ‘poor’ reliability between 13-22%, ‘fair’ reliability between 21-29% and ‘good’ to ‘excellent’ reliability between 49-66% (Figure 5a, Table 1). For the Desikan-Killiany cortical parcellation, connectomes reconstructed using DTI, CSD, GQI, GQI-DTI and CSD-DTI methods showed median ICC test-retest reliabilities between 0.52-0.66 and percentages of connections with ‘poor’ reliability between 15-32%, ‘fair’ reliability between 24-32% and ‘good’ to ‘excellent’ reliability between 37-61% (Figure 5b, Table 2). Test-retest reliability of the fractional anisotropy weighted connectomes showed similar results. Connectomes reconstructed using GQI-DTI and the Desikan-Killiany cortical parcellation showed a median ICC = 0.44 and ‘poor’ reliability in 42%, ‘fair’, reliability in 37% and ‘good’ or ‘excellent’ reliability in 21% of the connections (Figure 5, Table 3 and 4).

**Figure 5.**
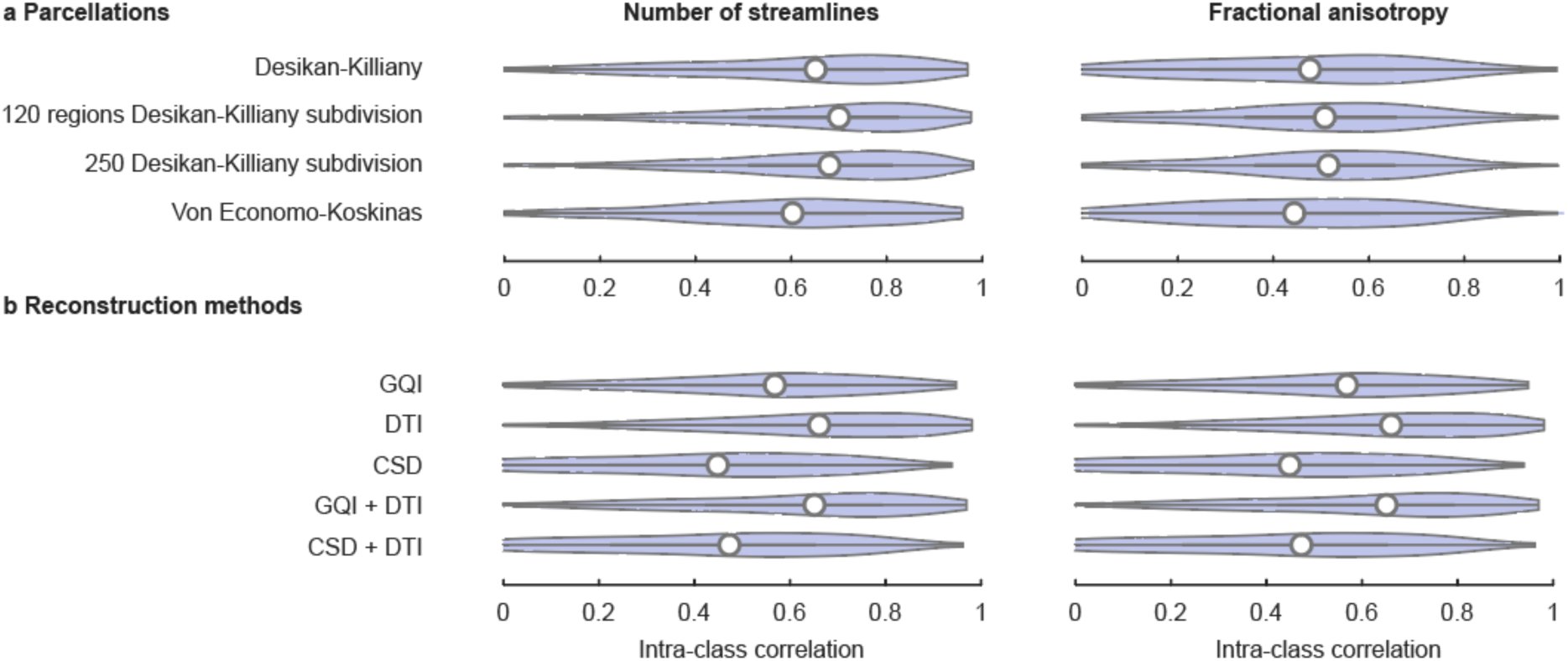
Structural connectome test-retest reliability. **a.** Test-retest reliability of structural connections for the Desikan-Killiany atlas, the 120 regions sub-parcellation of the Desikan-Killiany atlas, 250 regions sub-parcellation of the Desikan-Killiany atlas and the Von Economo-Koskinas atlas. **b.** Test-retest reliability of structural connections of the Desikan-Killiany atlas across different reconstruction methods. Boxes indicate the interval between the 25th and 75th percentiles (quartile *q*_1_ and *q*_3_), whiskers indicate the interval between *q*_1_ − 1.5 × (*q*_3_ - *q*_1_) and *q*_3_ + 1.5 × (*q*_3_-*q*_1_), the white circles indicate median values.

**Table 1.** Structural test-retest reliability across atlases weighted by number of streamlines.

| Reconstruction method | Median ICC | Interquartile range ICC | Percentage (%) of connections |  |  |
| --- | --- | --- | --- | --- | --- |
|  |  |  | Poor | Fair | Good or excellent |
| **Desikan-Killiany** | 0.66 | 0.47-0.79 | 19.5 | 23.6 | 57.0 |
| **120 regions Desikan-Killiany subdivision** | 0.70 | 0.54-0.82 | 13.2 | 21.8 | 65.0 |
| **250 regions Desikan-Killiany subdivision** | 0.69 | 0.54-0.81 | 13.0 | 21.1 | 65.9 |
| **Von Economo-Koskinas** | 0.59 | 0.42-0.75 | 22.3 | 29 | 48.8 |

**Table 2.** Structural test-retest reliability across methods weighted by number of streamlines.

| Reconstruction method | Median ICC | Interquartile range ICC | Percentage (%) of connections with reconstruction reliability |  |  |
| --- | --- | --- | --- | --- | --- |
|  |  |  | Poor | Fair | Good or excellent |
| **GQI** | 0.52 | 0.34-0.70 | 32.1 | 30.7 | 37.2 |
| **DTI** | 0.66 | 0.50-0.82 | 15.3 | 23.7 | 60.9 |
| **CSD** | 0.53 | 0.36-0.68 | 31.3 | 31.6 | 37.0 |
| **GQI + DTI** | 0.66 | 0.47-0.79 | 19.5 | 23.6 | 57.0 |
| **CSD + DTI** | 0.55 | 0.37-0.70 | 28.6 | 31.3 | 40.0 |

**Table 3.** Structural test-retest reliability across atlases weighted by fractional anisotropy.

| Reconstruction method | Median ICC | Interquartile range ICC | Percentage (%) of connections with reconstruction reliability |  |  |
| --- | --- | --- | --- | --- | --- |
|  |  |  | Poor | Fair | Good or excellent |
| Desikan-Killiany | 0.47 | 0.29-0.63 | 38.9 | 32.0 | 29.1 |
| 120 regions Desikan-Killiany subdivision | 0.51 | 0.36-0.64 | 30.8 | 36.8 | 32.4 |
| 250 regions Desikan-Killiany subdivision | 0.52 | 0.38-0.64 | 27.9 | 39.1 | 33.0 |
| Von Economo-Koskinas | 0.44 | 0.26-0.60 | 44.5 | 30.8 | 24.7 |

**Table 4.** Structural test-retest reliability across methods weighted by fractional anisotropy.

| Reconstruction method | Median ICC | Interquartile range ICC | Percentage (%) of connections |  |  |
| --- | --- | --- | --- | --- | --- |
|  |  |  | Poor | Fair | Good or excellent |
| GQI | 0.44 | 0.28-0.58 | 41.8 | 37.3 | 20.9 |
| DTI | 0.51 | 0.30-0.65 | 35.1 | 31.5 | 33.4 |
| CSD | 0.49 | 0.26-0.63 | 38 | 30.9 | 31.2 |
| GQI + DTI | 0.47 | 0.29-0.63 | 38.9 | 32 | 29.1 |
| CSD + DTI | 0.49 | 0.29-0.65 | 36.2 | 31 | 32.8 |

### Functional test-retest benchmarking

Test-retest reliability of the functional connectomes showed across atlases in 11-15% of the connections ‘poor’ reliability, 53-59% ‘fair’ reliability and in 26-36% ‘good’ to ‘excellent’ reliability (Figure 6a, Table 5). Focusing on the 68 cortical regions of the Desikan-Killiany parcellation, we observed that network reconstructions that included global mean regression resulted in 30% of the connections having ‘poor’, 49% ‘fair’ and 21% ‘good’ to ‘excellent’ reliability (median ICC=0.47, IQR=0.38-0.58). Leaving out the scrubbing preprocessing step (17% of the connections classified as having ‘poor’, 57% as having ‘fair’ and 27% as having ‘good’ to ‘excellent’ reliability,medianICC=0.52, IQR=0.44-0.61) or bandpass filter (11% of the connections classified as having ‘poor’, 58% ‘fair’ and 31% ‘good’ to ‘excellent’ reliability, median ICC=0.55, IQR=0.47-0.62) did not have a strong effect on the reconstruction reliability (Figure 6b, Table 6).

**Figure 6.**
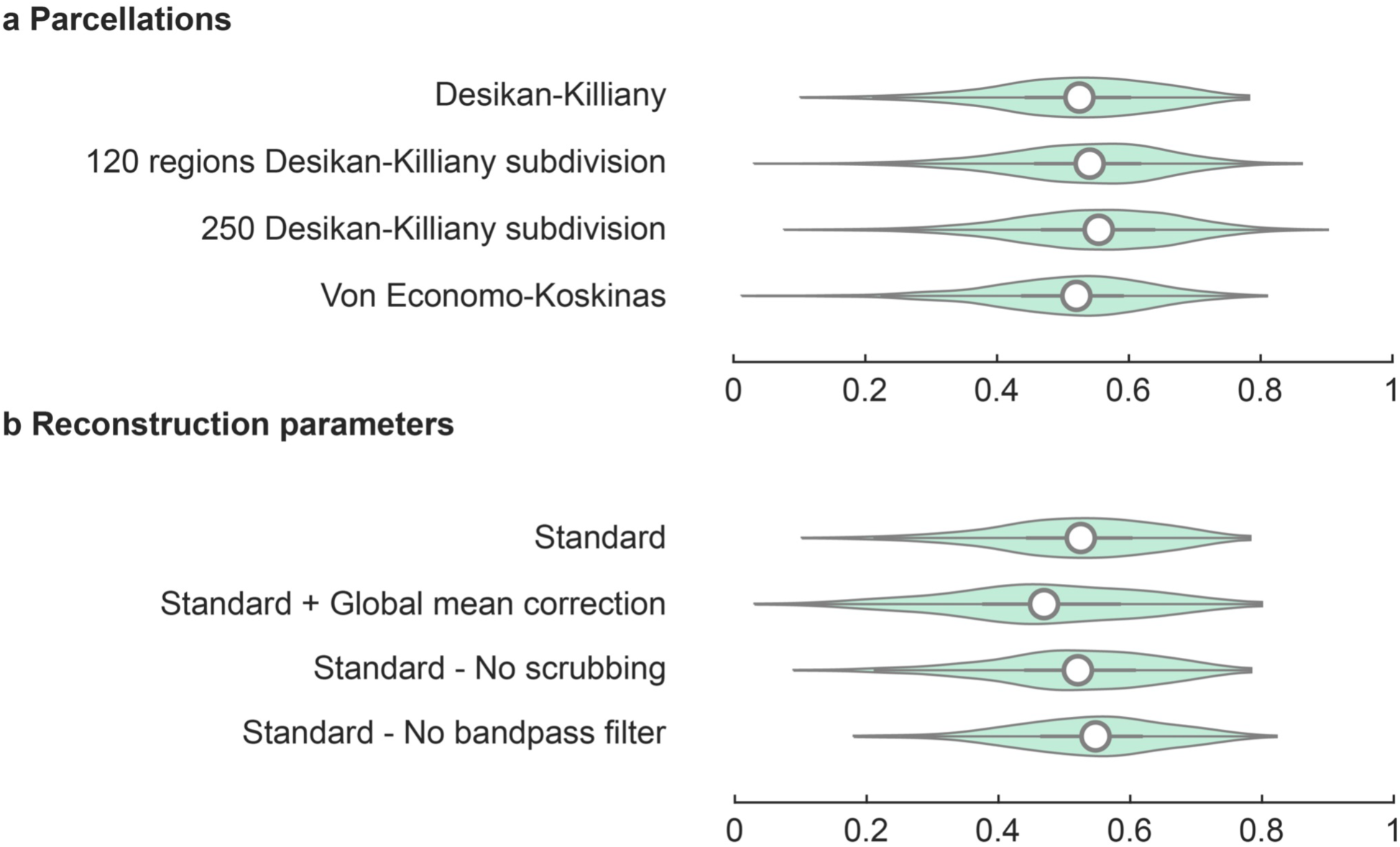
Functional pipeline test-retest reliability. **a.** Test-retest reliability of functional connections for the Desikan-Killiany atlas, the 120 regions sub-parcellation of the Desikan-Killiany atlas, 250 regions sub-parcellation of the Desikan-Killiany atlas and the Von Economo-Koskinas atlas. **b.** Test-retest reliability of functional connections of the Desikan-Killiany atlas, using standard HCP-default parameters (top), standard processing with additional global mean regression, standard processing excluding scrubbing and standard processing excluding bandpass filtering. Boxes indicate the interval between the 25th and 75th percentiles (quartile *q*_1_ and *q*_3_), whiskers indicate the interval between *q*_1_ − 1.5 × (*q*_3_ - *q*_1_) and *q*_3_ + 1.5 × (*q*_3_-*q*_1_), the white circles indicate median values.

**Table 5.** Functional test-retest reliability across atlases.

| Reconstruction method | Median ICC | Interquartile range ICC | Percentage (%) of connections with reconstruction reliability |  |  |
| --- | --- | --- | --- | --- | --- |
|  |  |  | Poor | Fair | Good or excellent |
| Desikan-Killiany | 0.52 | 0.44-0.60 | 15.1 | 58.8 | 26.1 |
| **120 regions Desikan-Killiany subdivision** | 0.54 | 0.46-0.62 | 13.4 | 56.8 | 29.8 |
| **250 regions Desikan-Killiany subdivision** | 0.55 | 0.47-0.64 | 11.6 | 52.7 | 35.7 |
| **Von Economo-Koskinas** | 0.53 | 0.46-0.60 | 10.9 | 55.5 | 21.9 |

**Table 6.** Functional test-retest reliability across methods.

| **Reconstruction method** | **Median ICC** | **Interquartile range ICC** | **Percentage (%) of connections with reconstruction reliability** |  |  |
| --- | --- | --- | --- | --- | --- |
|  |  |  | **Poor** | **Fair** | **Good or excellent** |
| **Standard** | 0.52 | 0.44-0.60 | 15.1 | 58.8 | 26.1 |
| **Standard + global mean regression** | 0.47 | 0.38-0.58 | 29.6 | 49.3 | 21.1 |
| **Standard - No scrubbing** | 0.52 | 0.44-0.61 | 16.7 | 56.6 | 26.8 |
| **Standard - No bandpass filtering** | 0.55 | 0.47-0.62 | 10.5 | 58.3 | 31.1 |

## Discussion

This software paper discusses CATO, a software toolbox for the reconstruction of structural and functional connectome maps. We describe the modular processing steps in the structural and functional pipelines and detail the implemented reconstruction methods, together with benchmark values to provide researchers insight in the sensitivity-specificity trade-off and reliability of connectome reconstructions.

Benchmarking structural reconstruction parameters and methods with respect to the ITC2015 challenge, shows that reconstruction methods in CATO have similar strengths and difficulties in reconstructing tracts as submissions to the ITC2015 challenge. ITC2015 datasets generally show difficulty in correctly reconstructing cortico-spinal and commissural tracts (see Figure 3 from (Maier-Hein et al., 2017)), and so does CATO. The anterior commissure and posterior commissure, tracts that were not reconstructed were considered “very hard” in the ITC2015 challenge, likely due to their small cross-sectional diameter (< 2mm) and the left cortico-spinal tract was considered “hard” to reconstruct. We believe that the percentage of valid fibers in CATO reconstructions could increase when reconstructed fiber clouds are further post-processed. Best performing DTI-based submissions of the ITC2015 challenge performed editing to clean up the fiber cloud. Such post-processing can be achieved manually, but also through semi-automatic methods such as fiber clustering in which fiber bundles are detected and small fiber clusters excluded (Garyfallidis et al., 2012) or by filtering out potential spurious connections based on anatomical priors (Schiavi et al., 2020).

Benchmarking the sensitivity and specificity of the reconstructed structural connectomes showed that implemented reconstruction methods, at their default settings, have relatively high specificity compared to submissions of the ITC2015 challenge (Figure 4). Such strict connectome reconstructions are beneficial as false positive connections are estimated to have more impact on network measures than false negatives (Zalesky et al., 2016). They do however have a negative effect on the level of sensitivity of finding more complex fiber bundles, i.e., at the price of having more false negatives (Helwegen et al., 2022). Exploring the effect of reconstruction parameters showed various ways in which researchers can tune the sensitivity and specificity profile of their reconstructions (Supplementary Figure 3). These benchmark findings suggest that the presented methods perform within the range of other powerful structural connectome reconstruction software tools and provide a competitive connectome reconstruction toolset with the flexibility to tune the sensitivity and specificity trade-off of reconstructions.

The HCP retest dataset provided the opportunity to estimate the test-retest reliability of connectomes processed. Structural connectivity showed higher reliability for the number of streamlines (NOS) connectivity strength (80% of the connections in the group matrix having ‘fair’ or better reliability) compared to fractional anisotropy connectivity strength (65%). Reconstructed functional connectivity showed reasonable reliability with around 70% of the connections in the group matrix having ‘fair’ or better reliability (i.e. ICC > 0.4). Median ICC scores showed around 0.40 which is on par with other functional methods that display an average ICC of individual connections of around 0.29 (Noble et al., 2019). We note that ICC measures only provide an estimate of the test-retest reliability; high ICC coefficients can also result from sensitivity to persistent covariates.

In addition to reconstruction accuracy, computational efficiency is an important factor when using connectome reconstruction software. CATO uses one computational thread by default and, for “clinical”-resolution data (2 mm isotropic voxels), up to 2 GB memory (this is specified in the `maxNumberCompThreads` and `maxMemoryGB` parameters). The duration of processing a subject depends most strongly on the data resolution and the preprocessing steps defined in the preprocessing scripts. To give an indication, structural processing takes approximately 30 minutes for preprocessed “clinical”-resolution (2 mm isotropic voxels) data and up to 2 hours for high-resolution (1.25 mm isotropic voxels) preprocessed HCP data. Functional processing takes between 15 minutes (clinical-resolution data) to 30 minutes (high-resolution data).

A number of remarks should be taken into account when interpreting the findings presented and when using CATO. First, methodological issues inherent to diffusion MRI and rs-fMRI have to be considered: diffusion MRI is limited by providing only an indirect measure of connectivity and no information on the directionality of connections (Mori and Zhang, 2006). Resting-state fMRI is limited by depending on correlations in the BOLD signal intensity (Constable, 2006). Both methods are further limited by their spatial and temporal resolution and can be sensitive to physiological motion (Constable, 2006; Mori and Zhang, 2006). Second, the user-provided preprocessing script requires attention to ensure that preprocessing steps do not interfere (Lindquist et al., 2019). Third, we focused in this paper on benchmarking connectome reconstructions, rather than optimizing performance. A recent study suggests that careful selection of network reconstruction parameters and preprocessing steps can improve the reconstruction quality and mitigate, for example, the impact of physical motion on structural connectivity (Oldham et al., 2020) and functional connectivity (Esteban et al., 2019). Fourth, the generalizability of benchmark results is inherently limited to the variety of datasets used in the benchmarking process. Here, the calibration of reconstruction methods and parameters was performed in two high-quality datasets. Future benchmarking in datasets with different acquisition protocols and data qualities is warranted to gain a better understanding of the performance of reconstruction methods and parameters.

The availability of a diverse set of connectivity reconstruction packages enables the neuroscience community to replicate reported results using multiple analysis pipelines to demonstrate their robustness to variations in applied processing software (Goodman et al., 2016). Earlier versions of CATO have been used in previous studies examining connectome organization in the healthy brain (van den Heuvel et al., 2019), across various diseases (e.g. (Repple et al., 2020)), neonatal brain organization (Keunen et al., 2018) and comparative connectomics (Ardesch et al., 2021). CATO is presented as an open-source connectivity reconstruction toolbox, shared under the MIT License.

## Acknowledgements

Siemon C. de Lange was supported by the Amsterdam Neuroscience alliance grant and ZonMw Open Competition, project REMOVE 09120011910032. Martijn P. van den Heuvel was supported by a VIDI (452-16-015) grant from the Netherlands Organization for Scientific Research (NWO) and Consolidator grant of the European Research Council (ERC CONNECT 101001062).

