## Supplementary figures and images for "Structural and functional connectivity reconstruction with CATO - A Connectivity Analysis TOolbox"

### Supplementary Figure 1

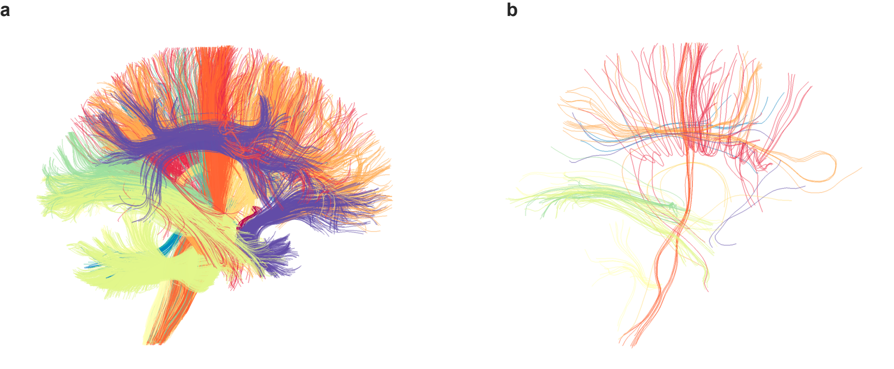

### Supplementary Figure 2

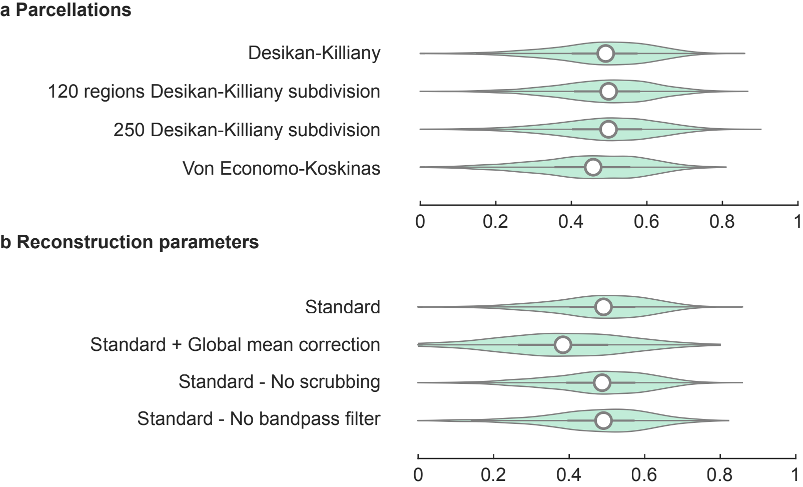

### Supplementary Figure 3

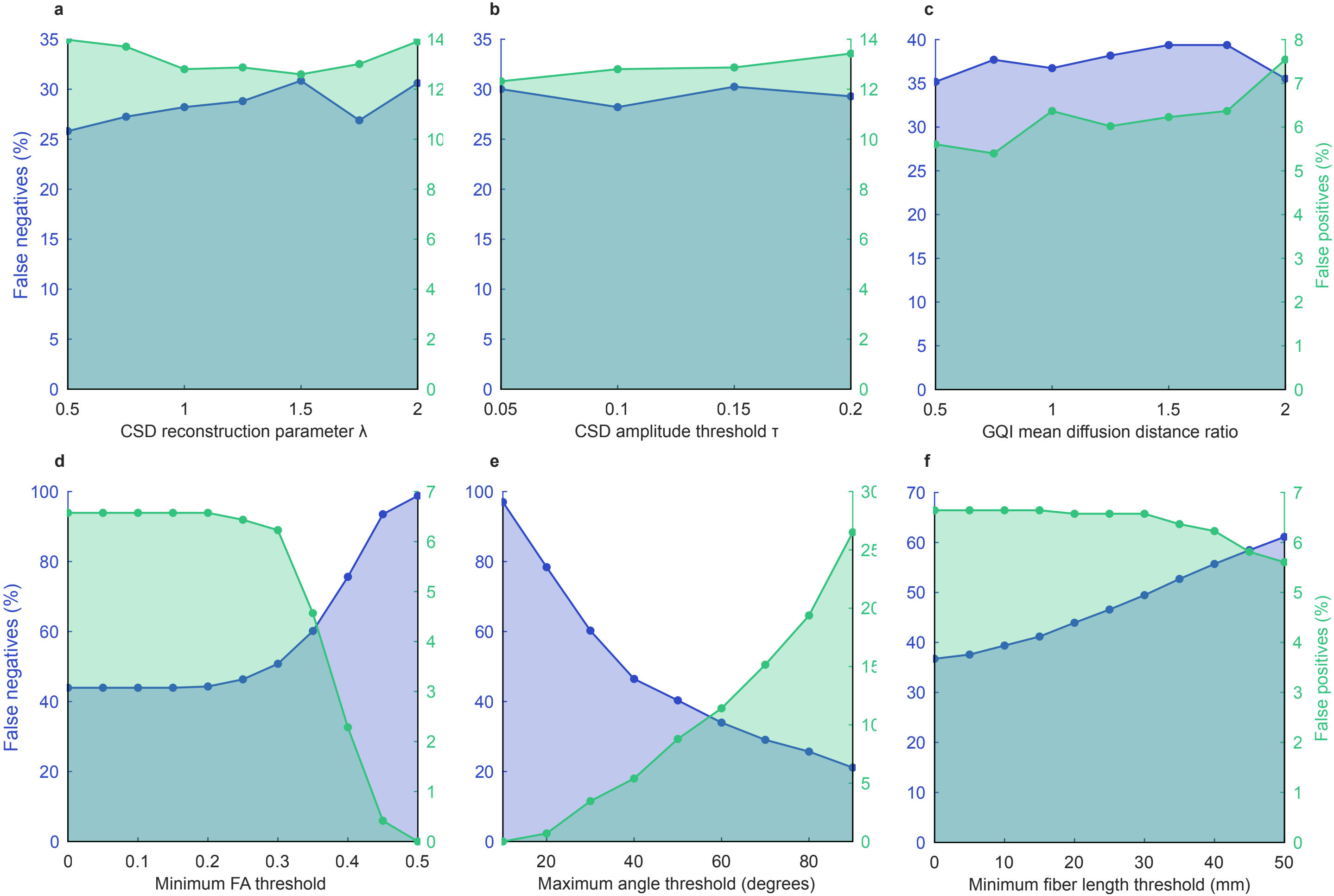
