## Supplementary Materials for "Structural and functional connectivity reconstruction with CATO - A Connectivity Analysis TOolbox"

### Supplementary Methods

#### Supplementary Methods 1 - Implementation iRESTORE algorithm

The iRESTORE diffusion reconstruction method identifies, per voxel, measurements that are outliers and that are excluded in the tensor fitting. To ensure that enough information is preserved for reliable tensor estimation, the iRESTORE algorithm checks that the B-matrix remains well conditioned and directionally balanced. When these requirements are not satisfied, then all measurements are used to perform the nonlinear least-squares fitting. The B-matrix is considered well-conditioned if the condition number is lower than `thresCondNum` and directionally balanced if the variation in average projection scores is lower than `thresVarProjScores`. The condition number and variation in average projection scores thresholds are specific for each gradient acquisition scheme and are estimated by the function `thresholdAssistant`. Following de Reus (de Reus, 2015), each threshold is estimated as the average of two threshold estimates. Each threshold estimate is obtained from bootstrapping a sample of gradient schemes in which random gradient directions are removed from the original scheme (1,000 permutations). The first threshold estimate is the value such that the removal of 5% of the gradients is accepted in 75% of the samples. The second threshold estimate is the value such that the removal of 50% of the gradients is accepted in 25% of the samples. The suggested threshold for the condition number and variation in average projection scores is then the average of these two estimates.

The non-linear least squares fitting procedure used parameters suggested by Gavin et al. (Gavin, 2019) ($\epsilon_1 = 0.001$, $\epsilon_2 = 0.001$, $\epsilon_3 = 0.1$, $\epsilon_4 = 0.1$, $\lambda_\uparrow = 11$, $\lambda_\downarrow = 9$, $\lambda_0 = 0.01$ and a maximum of 100 iterations).

**Supplementary Methods 2 - ITC2015 CATO reconstruction**

Whole brain segmentation was performed on the anatomical T1-weighted image using FreeSurfer version 6.0.0 (Fischl et al., 2004). The example topup-eddy-preprocessing script provided with CATO was used to correct DWI images for motion, susceptibility induced distortions and eddy current distortions using FSL version 5.0.10 (Jenkinson et al., 2012). White matter pathways were reconstructed using diffusion reconstruction methods DTI, CSD, GQI and GQI-DTI. In the DTI reconstruction method, the threshold on the condition number for selecting non-outlying measurements and the threshold on the variation in the average projection scores for selecting non-outlying measurements were automatically estimated by CATO. The following parameters were used in the GQI reconstruction method: mean diffusion distance ratio = 1.25, number of maximum output peaks = 4, minimum peak coefficient below which peaks are discarded = 0, maximum number of identified peaks beyond which a voxel is considered isotropic = 8. The CSD reconstruction was performed with the following parameters: $\lambda = 1.25$, spherical harmonics order = 10 and $\tau = 0.1$, number of maximum output peaks = 4, minimum peak coefficient below which peaks are discarded = 0.1, maximum number of identified peaks beyond which a voxel is considered isotropic = 8).

Following the ITC2015 instructions, fiber tracing started from voxels in the white-matter, brainstem and cerebellum (8 seeds per voxel were used). Fiber tracing followed the main diffusion direction of each voxel and propagated until a streamline reached one of the stopping criteria (i.e. the fiber was about to exit the brain mask, made a turn of > 45 degrees and/or reached a voxel with low fractional anisotropy < 0.1).

In addition to benchmarking the four diffusion reconstruction methods with default parameter settings, the effects of specific reconstruction parameters on the sensitivity and specificity were investigated. Connectomes were reconstructed with CSD reconstruction parameter $\lambda$ spanning the range [0.5 - 2], CSD amplitude thresholds $\tau$ in the range [0.05 - 0.2] and GQI mean diffusion distance ratios in the range [0.5 - 2]. To understand the effect of the fiber tracking parameters, connectomes were constructed for minimum FA thresholds in the range [0 - 0.5], maximum angle thresholds in the range [0 - 90] degrees and minimum fiber length thresholds in the range [0 - 50] mm. The explored threshold varied while the other parameters were set to minimum FA threshold = 0.1, maximum angle threshold = 45° or minimum fiber length threshold = 2 cm.

Structural connectivity reconstruction for the ITC2015 challenge was performed on an iMac (Late 2015) with macOS Mojave 10.14.6 as operating system and with a 3.2 GHz Intel Core i5 processor with 16 GB memory.

**Supplementary Methods 3 - HCP connectivity reconstruction**

The HCP provided cortical surface reconstructions and brain segmentations obtained using FreeSurfer (version 5.3-HCP). Using FreeSurfer additional cortical surface annotations were constructed for the 120, 250 and 500 regions Cammoun sub-parcellations of the Desikan-Killiany atlas and the Von Economo-Koskinas cortical region and cortical-type atlas. We further ensured that voxels in the cortical parcellations were mapped to the correct hemisphere indicated by the hemisphere assignment in the brain segmentation file.

Two rs-fMRI datasets, from different sessions during the same imaging visit, were used for each subject and were processed separately. A custom preprocessing script was used that calculated motion parameters from the movement regression file provided by the HCP and mapped the segmentation file to rs-fMRI reference image. Network reconstruction was performed with default HCP reconstruction parameters (van den Heuvel et al., 2017): 1. The signal was corrected for global effects by regressing out effects of motion (using the six motion parameters) and signal in CSF and left and right white matter; 2. The signal was band-pass filtered between 0.01 and 0.1 Hz; The signal was scrubbed by removing frames with potential movement artifacts as indicated by a framewise displacement larger than 0.25 and a DVARS value 1.5×IQR above the third quartile. Moreover, 1 frame preceding each frame with potential movement artifacts was also removed to accommodate temporal smoothing of the signal.

Structural test-retest benchmarking was performed on DWI data from the test and retest visits. White matter pathways were reconstructed using diffusion reconstruction methods DTI, CSD, GQI, CSD-DTI and GQI-DTI. Diffusion reconstruction using DTI was performed with automatically estimated reconstruction parameters. For the GQI reconstruction method the following parameters were used: mean diffusion distance ratio = 1.25, number of maximum output peaks = 4, minimum peak coefficient below which peaks are discarded = 0, maximum number of identified peaks beyond which a voxel is considered isotropic = 8. CSD diffusion reconstruction was performed with the following parameters: $\lambda = 1$, spherical harmonics order = 6 and $\tau = 0.1$, number of maximum output peaks = 4, minimum peak coefficient below which peaks are discarded = 0, maximum number of identified peaks beyond which a voxel is considered isotropic = 8). Fiber tracing started from voxels in the white matter (8 seeds per voxel). Fiber tracing followed the main diffusion direction of each voxel and propagated until a streamline reached one of the stopping criteria (i.e. the fiber was about to exit the white and gray matter, made a > 45 degrees turn and/or reached a voxel with low fractional anisotropy < 0.1).

Structural and functional reconstruction was performed on a Mac Pro (Mid 2010) with macOS Sierra 10.12.6 as operating system and with a 2 x 2,66 GHz 6-Core Intel Xeon processor with 64 GB memory.

### Supplementary Results

#### Supplementary Results 1 - Connectome reconstruction parameters

CATO reconstruction parameters can be used to tune the sensitivity and specificity of connectome reconstructions. Benchmarking CSD reconstruction parameter λ (regulating the coarseness of the reconstructed peak profile), showed that higher values of λ result in higher reconstruction specificity (λ = 0.5 FN: 26% and FP: 14%; and λ = 2 FN: 31% and FP: 14%) (Supplementary Figure 3a), but this effect was small (<5%) compared to the variance in FN and FP across reconstruction methods (Figure 4c). Changing CSD amplitude threshold τ showed also no large (>5%) effect on the false negative and false positive rates (τ = 0.05 FN: 30% and FP: 12%; and τ = 0.2 FN: 29% and FP: 13%, Supplementary Figure 3b). Mean diffusion distance ratio parameter (MDDR) regulates the coarseness of the peak profile in the GQI reconstruction method and benchmarking the mean diffusion distance ratio showed a complex pattern in which the specificity decreased from MDDR = 0.5 (FN: 35% and FP: 6%) to MDDR = 2.0 (FN: 36% and FP: 8%) (Supplementary Figure 3c). These benchmark results suggest that, in the explored parameter range, diffusion reconstruction parameters show small effects on the false negatives and false positives rates compared to the differences across reconstruction methods (i.e. DTI versus CSD).

To further evaluate the effect of the minimum FA, minimum fiber length and maximum turn angle thresholds, we reconstructed fiber clouds in which each time one of these thresholds was varied. The minimum FA threshold in the fiber reconstruction step tuned the connectome between highly sensitive reconstructions when no FA threshold was applied (FA threshold = 0, FN: 44%, FP: 7%) to highly specific results when strict FA thresholds were applied (e.g. FA threshold = 0.5, FN:99%, FP: 0%) (Supplementary Figure 3d). The maximum angle threshold (MAT) tuned connectome reconstructions in a wide range of specificity-sensitivity trade offs (Supplementary Figure 3e). Applying no threshold on the maximum angle resulted in a lean connectome reconstruction with a relative high number of false positives (MAT = 90º FN: 21%, FP: 27%) and the most stringent threshold (MAT = 10º) resulted in the most specific reconstruction possible with no reconstructed connections (FN: 97% and FP: 0%). Applying no minimum fiber length threshold resulted in the most sensitive reconstructions (FN: 37%, FP: 7%) and filtering fibers above 50 mm resulted in connectomes with more specificity (FN: 61%, FP: 6%) (Supplementary Figure 3f). These results showed that settings for the fiber reconstruction thresholds have a large effect on the connectome reconstruction quality and sensitivity / specificity balance and are meaningful reconstruction parameters for researchers to explore in studies.

### Supplementary Figures

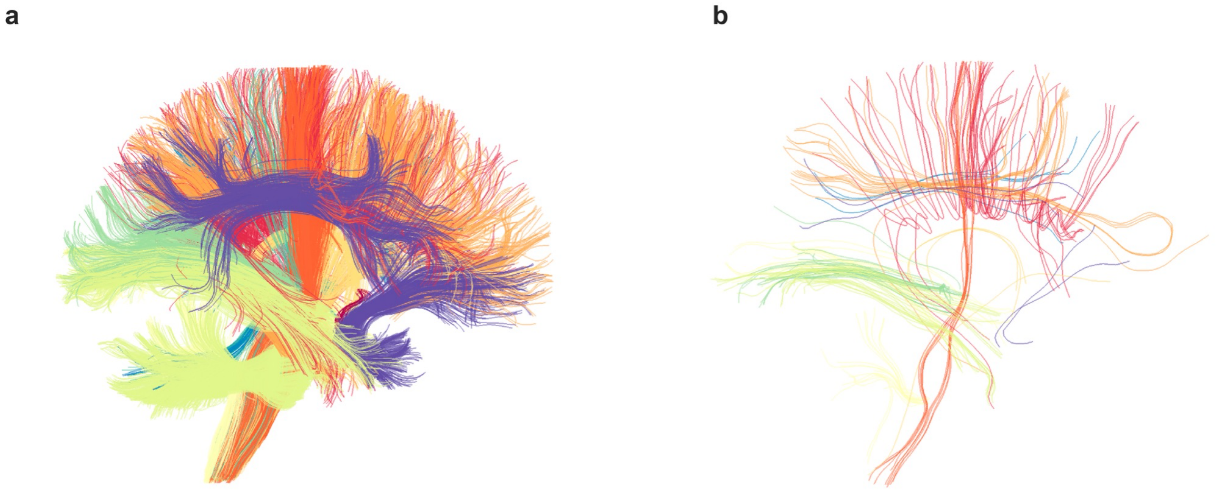


**Supplementary Figure 1. Ground truth fibers included in the ground truth connectivity matrix. a.** Ground truth fibers included in the ground truth connectivity matrix. **b.** Fibers from the ground truth fiber cloud not included in the ground-truth connectivity matrix. On average 96.4% of the fibers in the ground truth fiber cloud were also included in a connectivity matrix (that included cortical regions, subcortical regions, cerebellum and brain stem as nodes). Most white matter bundles showed very high >99 % (11/25 bundles) or high percentages > 95% (21/25 bundles). For visualization purposes, the figure shows a sample of 2% of the fibers from each fiber cloud.

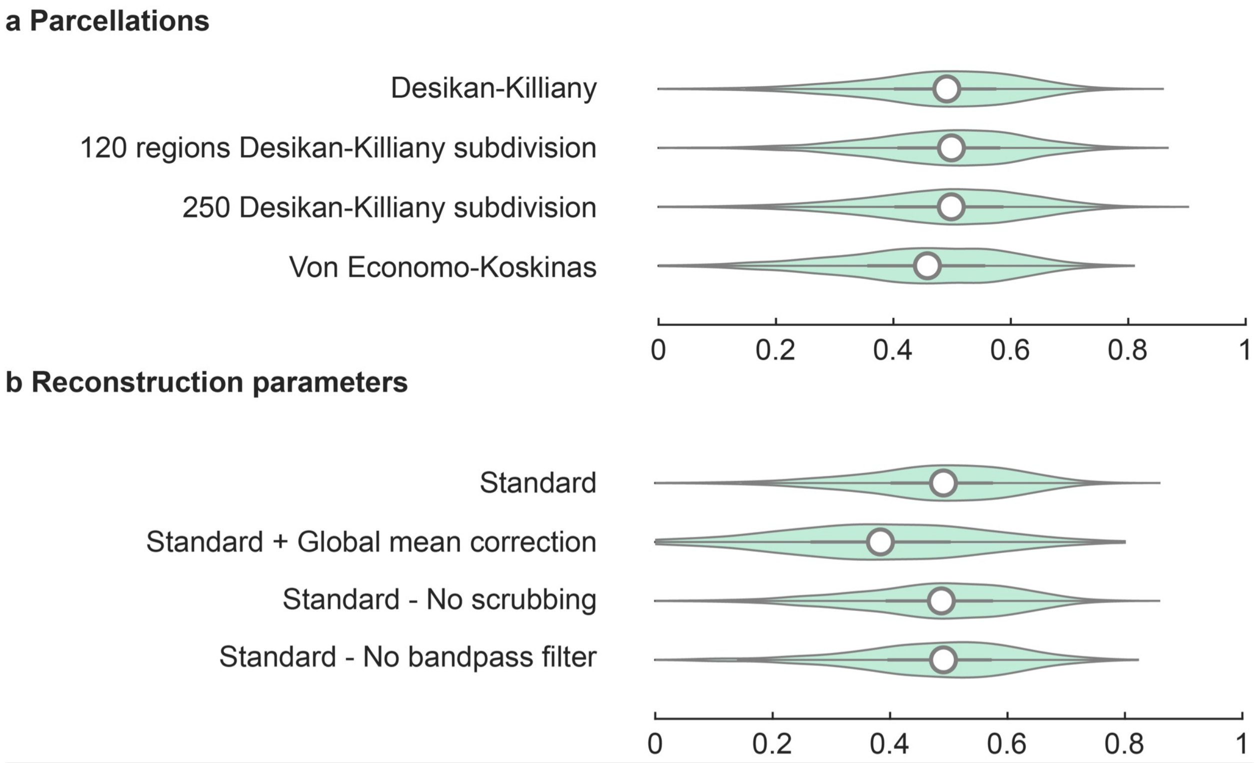


**Supplementary Figure 2. Functional pipeline test-retest reliability across all connections. a.** Test-retest reliability of functional connections (including connections not in the group-matrix) for the Desikan-Killiany atlas, the 120 regions sub-parcellation of the Desikan-Killiany atlas, 250 regions sub-parcellation of the Desikan-Killiany atlas and the Von Economo-Koskinas atlas. **b.** Test-retest reliability of functional connections of the Desikan-Killiany atlas, using standard HCP-default parameters (top), standard processing with additional global mean regression, standard processing excluding scrubbing and standard processing excluding bandpass filtering. Boxes indicate the interval between the 25th and 75th percentiles (quartile $q_1$ and $q_3$), whiskers indicate the interval between $q_1 - 1.5 \times (q_3 - q_1)$ and $q_3 + 1.5 \times (q_3 - q_1)$, the white circles indicate median values.

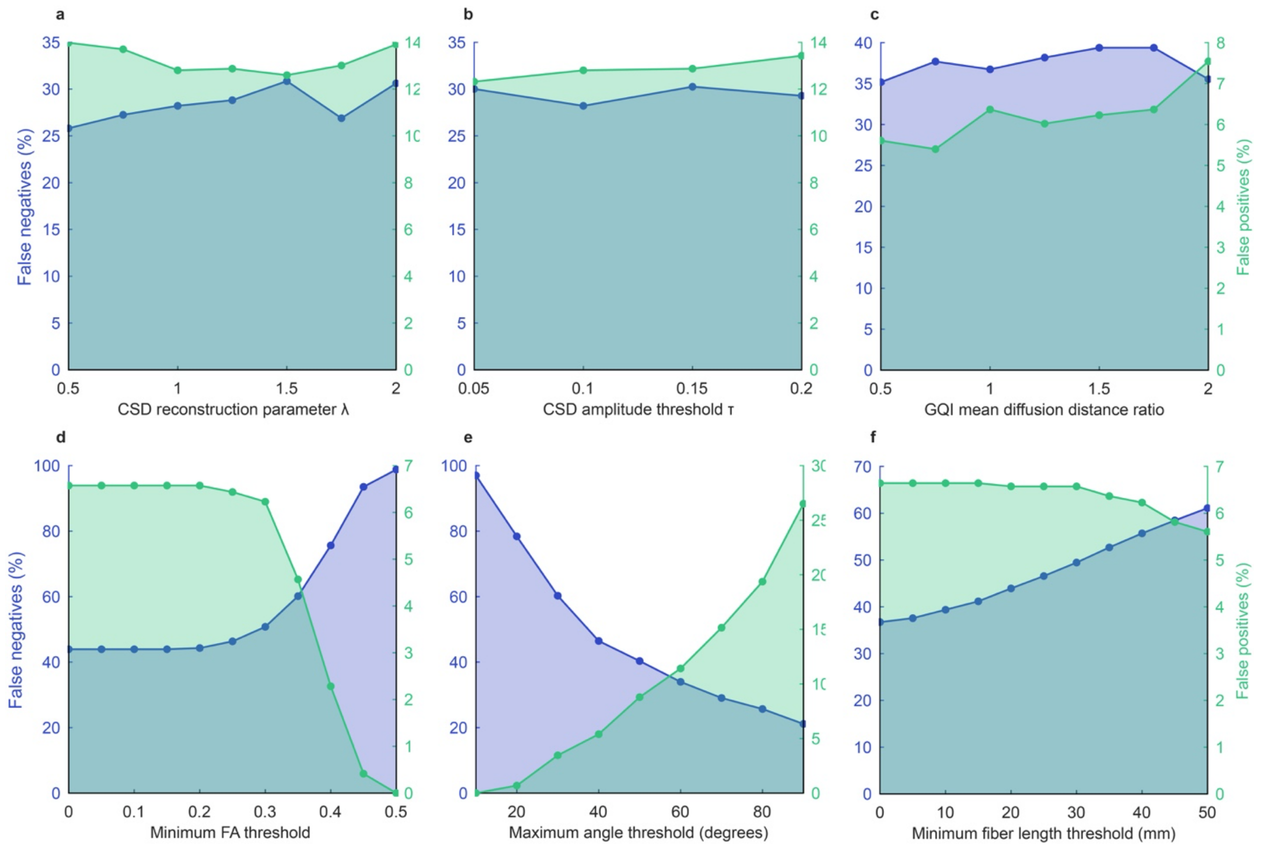


**Supplementary Figure 3. Effect of reconstruction settings on connectome reconstruction sensitivity and specificity.** The effect of CSD diffusion reconstruction parameters λ and τ and GQI mean diffusion distance ratio on the percentage of false positives and false negatives was examined on the first row (**a**, **b** and **c**). These diffusion reconstruction parameters showed in the explored range small effects on the sensitivity and specificity of the reconstructed connectomes compared to the differences observed between reconstruction methods. The second row shows the effect of minimum fractional anisotropy (FA) threshold, maximum turn angle threshold and minimum fiber length threshold (**d**, **e** and **f**). These fiber reconstruction thresholds showed a major impact on the sensitivity and specificity of the reconstructed connectivity matrices. Increasing the minimum fiber length and minimum FA threshold increased the specificity, whereas higher maximum angle thresholds resulted in more lenient reconstructions with higher sensitivity.

### Supplementary Tables

#### Supplementary Table 1. Overview of white matter bundles in ground truth white matter skeleton in the ISMRM 2015 Tractography challenge.

| Abbreviation | Full name |
|---|---|
| CA | Anterior commissure |
| CC | Corpus callosum |
| CG - left | Cingulum - left |
| CG - right | Cingulum - right |
| CP | Posterior commissure |
| CST - left | Cortico-spinal tract - left |
| CST - right | Cortico-spinal tract - right |
| Fornix | Fornix |
| FPT - left | Frontopontine tracts - left |
| FPT - right | Frontopontine tracts - right |
| ICP - left | Inferior cerebellar peduncle - left |
| ICP - right | Inferior cerebellar peduncle - right |
| ILF - left | Inferior longitudinal fasciculus - left |
| ILF - right | Inferior longitudinal fasciculus - right |
| MCP | Middle cerebellar peduncle |
| OR - left | Optic radiation - left |
| OR - right | Optic radiation - right |
| POPT - left | parieto-occipital pontine tract - left |
| POPT - right | parieto-occipital pontine tract - right |
| SCP - left | Superior cerebellar peduncle - left |
| SCP - right | Superior cerebellar peduncle - right |
| SLF - left | Superior longitudinal fasciculus - left |

| | |
|---|---|
| SLF - right | Superior longitudinal fasciculus - right |
| UF - left | Uncinate fasciculus - left |
| UF - right | Uncinate fasciculus - right |

**Supplementary Table 2. Functional test-retest reliability of all connections across atlases.**

| Reconstruction method | Median ICC | Interquartile range ICC | Percentage (%) of connections with reconstruction reliability | | |
|---|---|---|---|---|---|
| | | | Poor | Fair | Good or excellent |
| **Desikan-Killiany** | 0.49 | 0.40-0.57 | 24.5 | 57.6 | 18.0 |
| **120 regions Desikan-Killiany subdivision** | 0.50 | 0.41-0.58 | 23.3 | 57.2 | 19.5 |
| **250 regions Desikan-Killiany subdivision** | 0.50 | 0.40-0.59 | 24.3 | 54.3 | 21.4 |
| **Von Economo-Koskinas** | 0.46 | 0.36-0.55 | 34.3 | 52 | 13.8 |

**Supplementary Table 3. Functional test-retest reliability of all connections across methods.**

| Reconstruction method | Median ICC | Interquartile range ICC | Percentage (%) of connections with reconstruction reliability | | |
|---|---|---|---|---|---|
| | | | Poor | Fair | Good or excellent |
| **Standard** | 0.49 | 0.40-0.57 | 24.5 | 57.6 | 18.0 |
| **Standard + global mean regression** | 0.38 | 0.27-0.50 | 53.6 | 36.9 | 9.5 |
| **Standard - No scrubbing** | 0.49 | 0.39-0.57 | 26.2 | 55.8 | 18.1 |
| **Standard - No bandpass filtering** | 0.49 | 0.40-0.57 | 25.7 | 56.5 | 17.8 |
